# Structural and Functional Studies of Rabbit SAMD9 Reveal a Distinct tRNase Module That Underlies the Antiviral Activity

**DOI:** 10.1101/2025.04.10.648150

**Authors:** Juhi Chaturvedi, Fushun Zhang, Chen Zhang, Sonal Badhe, Yan Xiang, Junpeng Deng

## Abstract

Human SAMD9 and SAMD9L (collectively SAMD9/9L) are large cytoplasmic proteins with antiviral and antiproliferative activities, recently shown to regulate protein synthesis by specifically cleaving phenylalanine tRNA (tRNA^Phe^). The enzymatic activity of human SAMD9 (hSAMD9) resides within its N-terminal tRNase domain, which depends on three essential basic residues for tRNA binding and biological activity. While these residues are highly conserved across mammalian SAMD9/9L, lagomorph SAMD9 orthologs uniquely harbor a charge-reversal acidic residue at one of three sites, a change known to inactivate hSAMD9/9L. Here, we show that despite this variation, rabbit SAMD9 (rSAMD9) potently restricts vaccinia virus replication and specifically reduces tRNA^Phe^ levels, mirroring hSAMD9. However, unlike hSAMD9, rSAMD9’s minimal tRNase module extends beyond the homologous tRNase domain (amino acid 158-389) to include the SIR2 region. Additional basic residues, one unique to rSAMD9, were also found to be important for its antiviral activity. The crystal structure of rSAMD9^158–389^ closely resembles hSAMD9^156–385^, though with difference in loop conformations. These findings demonstrate that lagomorph SAMD9 preserves core tRNA-targeting and antiviral functions despite a key residue variation and the need for an extended tRNase module.

**AUTHOR SUMMARY:** Sterile Alpha Motif Domain-containing 9 (SAMD9) and its paralog SAMD9-like (SAMD9L) are antiviral factors and tumor suppressors. Gain of function (GoF) mutations of SAMD9/9L, however, cause a spectrum of human diseases with immunological or/and neurological presentations. We recently identified the effector domain of SAMD9/9L as a previously unrecognized tRNase, a function critical for both their normal physiological roles and the pathogenic effects of by patient-derived mutations. We also identified three basic residues within the tRNase domain that are highly conserved across mammalian SAMD9/9L as key to its enzymatic and biological activities. However, lagomorph SAMD9 orthologs uniquely harbor a negatively charged acidic residue at one of three sites. Through extensive structural and functional analyses, here we demonstrate that rabbit SAMD9 (rSAMD9) retains both tRNA-cleaving and antiviral activities, despite notable differences in its tRNase module- including a unique basic residue specific to rSAMD9 and a larger domain architecture extending beyond the conserved tRNase core. These data highlight both the evolutionary conservation of tRNase activity in mammalian SAMD9/9L and the flexibility of the tRNase module within the protein family.

## Introduction

SAMD9 and SAMD9L are paralogous genes widely distributed in mammals[1]. In humans, they are tandemly located on chromosome 7 and share approximately 60% amino acid sequence identity. Both proteins (collectively referred to as SAMD9/9L) exhibit antiviral and antiproliferative functions[2-4], which have recently been linked to their ability to regulate cellular protein synthesis[5,6]. SAMD9/9L serve as key restriction factors against poxviruses [2- 4]. In turn, many mammalian poxviruses encode specific antagonists of SAMD9/9L. For instance, Vaccinia virus (VACV) expresses two such inhibitors, K1 and C7, each targeting distinct regions of SAMD9/9L [7,8]. Deletion of K1 and C7 abolishes VACV replication in most mammalian cell types [9-11].

Beyond their roles in viral restriction, heterozygous mutations of SAMD9/9L underlie several human developmental and hematologic diseases. These include MIRAGE syndrome [12], a multisystem disorder characterized by myelodysplasia, infection, restricted growth, adrenal hypoplasia, genital abnormalities, and enteropathy, and 10–20% of inherited bone marrow failure (IBMF) syndromes or pediatric myelodysplastic syndromes (MDS) [13-16]. In addition, SAMD9L mutations also cause ataxia-pancytopenia (ATXPC) syndrome, characterized by progressive neurological decline and bone marrow hypoplasia [17,18]. These diverse disorders are associated with gain-of-function (GoF) SAMD9/9L mutations that enhance the growth-suppressive activities of SAMD9/9L in vitro [12,15].

SAMD9/9L are large cytoplasmic proteins with more than 1,500 amino acids. They harbor multiple predicted domains from N- to the C-terminus: SAM (Sterile Alpha Motif), Alba (Acetylation Lowers Binding Affinity), SIR2 (Silent Information Regulator 2), P-loop NTPase, TPR (Tetratricopeptide Repeat), and OB (Oligonucleotide/Oligosaccharide Binding)[1]. This architecture resembles Nod-like receptors (NLRs), which consist of an N-terminal effector, a central nucleotide-mediated oligomerization domain, and a C-terminal sensor[19,20]. Recent studies of human SAMD9 (hSAMD9) identified a crucial N-terminal effector domain (amino acids 156–385), termed tRNase-SA, which serves as a tRNA endoribonuclease specifically cleaving tRNA^Phe^ [5,21]. SAMD9’s tRNase activity in cells can be activated by poxvirus infection or by patient-derived GoF mutations [21]. Once activated, SAMD9 depletes cellular tRNA^Phe^, causing codon-specific ribosomal stalling, inhibiting protein synthesis, and inducing proteotoxic stress.

Structural studies of hSAMD9’s tRNase-SA domain in complex with DNA has identified three basic residues, K198, K214, and R221, as essential for nucleic acid binding [5,21]. Charge-reversal mutations at any one of three key residues abolish hSAMD9’s antiviral and antiproliferative functions, and mutating the residue equivalent to R221 in human SAMD9L similarly disrupts its activity. Notably, lagomorph SAMD9 orthologs diverge from other mammalian SAMD9/9L by having an oppositely charged residue at the position corresponding to hSAMD9’s R221, raising questions about their functionality. Here, we present structural and functional studies of rabbit SAMD9, revealing that lagomorph SAMD9 proteins retain the core functions of hSAMD9 by utilizing a larger tRNase module and an alternative basic residue.

## Results

### Lagomorph SAMD9 proteins are unique among mammalian SAMD9/9L in having an oppositely charged residue at the R221-equivalent site

To assess whether the tRNase function is conserved across mammalian SAMD9/9L, we performed a phylogenetic analysis using approximately 370 mammalian SAMD9/9L proteins from the NCBI reference protein database and aligned their tRNase domain sequences (Figure 1 and S1). Previous structural and functional studies of hSAMD9 tRNase identified four acidic residues and three basic residues as essential for catalysis and tRNA binding, respectively[5,21]. Among the four acidic residues, E184, E196, and E218 are strictly conserved in every mammalian SAMD9/9L examined, while D241 is also conserved in most lineages but is replaced by an asparagine in SAMD9L from horses and many carnivores (Figure 1 and S1). Among the three basic residues, K214 is fully conserved; K198 is present in all but gray squirrel SAMD9L, which instead has a glutamate; and R221 is preserved across all lineages except for the four lagomorph SAMD9 sequences, which all have a glutamate at the R221-equivalent position (Figure 1). Two less critical basic residues (K350 and K242) were also identified in hSAMD9[5], and both are not strictly conserved and exhibit a greater number of substitutions in different taxa. For example, lagomorph SAMD9 proteins have a valine at the position corresponding to K350 (Figure 1). Hence, lagomorph SAMD9 uniquely retain only two of the three essential basic residues and one of the two less critical basic residues. By contrast, lagomorph SAMD9L preserves all of the critical acidic and basic residues.

**Figure 1.**
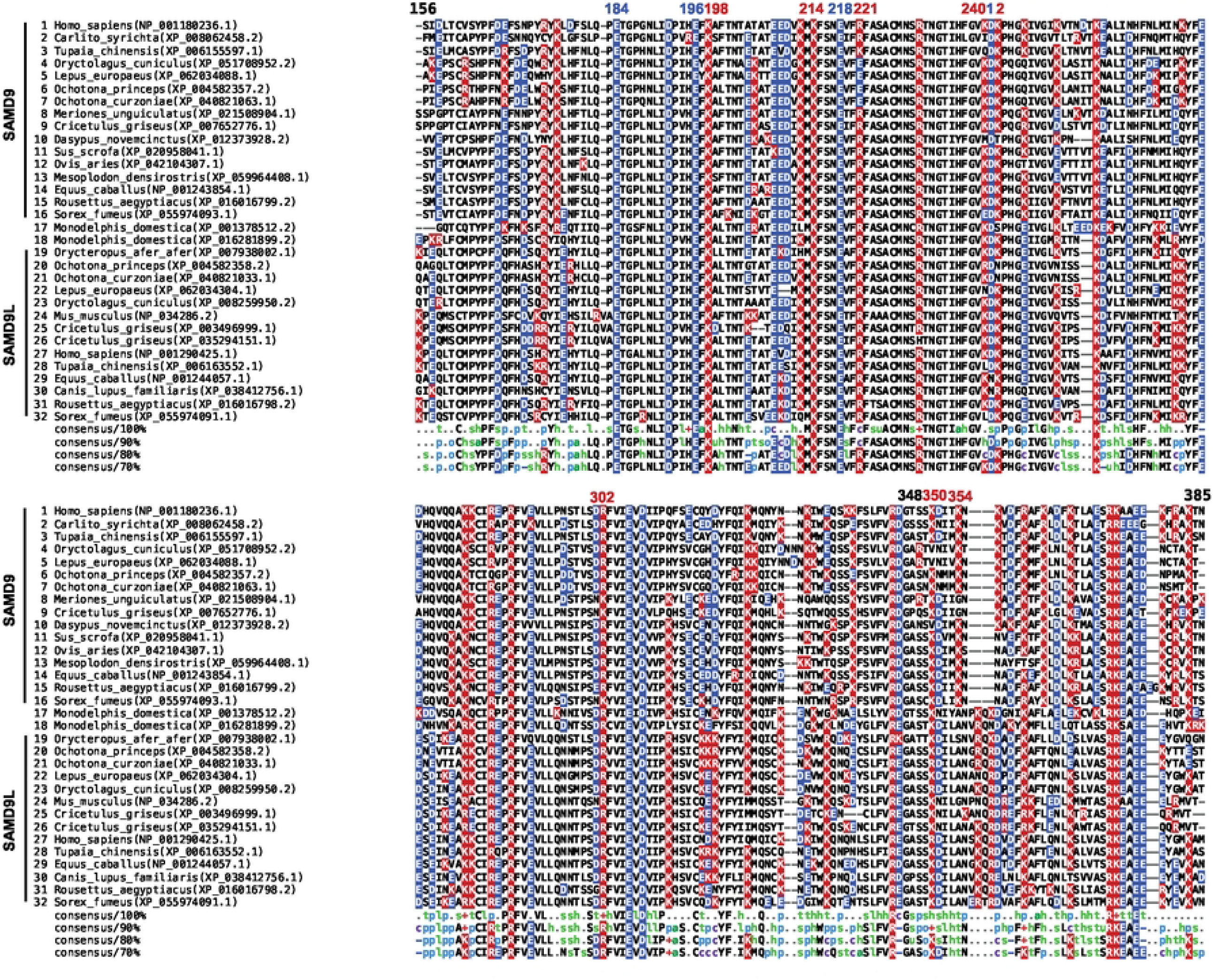
Multiple sequence alignment of the tRNase domain from representative mammalian SAMD9/9L proteins. Acidic (blue) and basic (red) residues are highlighted. Species names and protein accession numbers are shown alongside each sequence, with SAMD9 and SAMD9L entries clustered separately, except for two opossum (marsupial) SAMD9-like proteins that cannot be unambiguously assigned to either cluster. The numbering above the alignment indicates positions in human SAMD9.

### Rabbit SAMD9 exhibits antiviral activity comparable to human SAMD9

Rabbit SAMD9 (rSAMD9) was chosen as a representative lagomorph SAMD9 for functional analysis, since its gene can be readily cloned from the RK-13 rabbit cell line. Sequencing of the cDNA clone showed that it is nearly identical to the rabbit reference sequence. To evaluate rSAMD9 function, we tested the ability of rSAMD9 to inhibit vaccinia virus replication in HEK 293T cells, which does not express sufficient levels of endogenous human SAMD9/9L to restrict vaccinia virus [5]. An mCherry-tagged rSAMD9 plasmid was transfected into 293T cells, followed by infection with a vaccinia virus lacking viral SAMD9/9L inhibitors and encoding a GFP reporter (vK1^-^C7^-^/GFP^+^)[5]. SAMD9 and GFP expression levels were measured at the single-cell level via flow cytometry. Antiviral activity was quantified by comparing GFP expression in mCherry-positive cells with that in non-transfected cells in the same well. Wild type (WT) rSAMD9 inhibited viral GFP expression, exhibiting robust antiviral activity similar to that of hSAMD9 (Figure 2).

**Figure 2.**
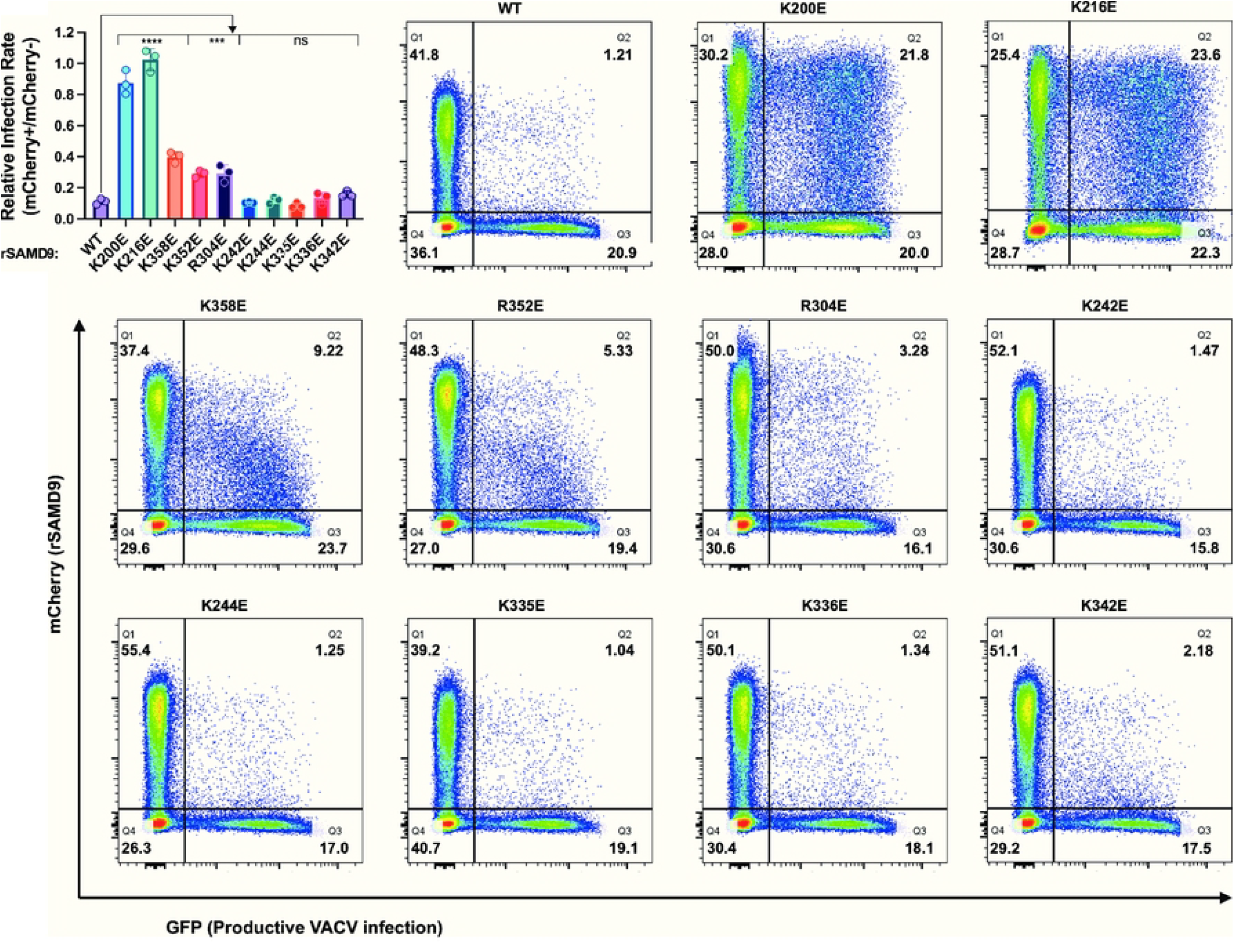
Rabbit SAMD9 requires specific basic residues in its tRNase domain for antiviral activity. HEK 293T cells were transfected with mCherry-rSAMD9 (wild-type or the indicated mutations) and infected with vK1^-^C7^-^/GFP^+^. Representative flow cytometry plots from one of the biological replicates are shown, with percentage of cells in each quadrant indicated. Relative infection rates between rSAMD9-expressing and nontransfected cells are derived from the flow cytometry data. Each biological replicate and SD are shown. Statistics: one-way ANOVA compared to the WT (ns, not significant; ****P < 0.0001; ***P < 0.001).

To determine whether the putative tRNase domain is essential for this antiviral activity, we performed charge-reversal mutations of the putative tRNA binding residues in rSAMD9. Mutations of K200 or K216, corresponding to hSAMD9 K198 and K214, abolished antiviral activity, allowing comparable levels of viral replication in cells regardless of SAMD9 expression. This result mirrors previous observations for hSAMD9. By contrast, mutating K244, which corresponds to hSAMD9 K242, did not reduce antiviral activity (Figure 2).

To assess whether basic residues adjacent to the putative tRNA-binding surface could compensate for the lack of R221- and K350-equivalent residues, we performed additional charge reversal mutations. While rSAMD9 with K242E, K335E, K336E, or K342E substitutions retained full antiviral activity, variants with R304E, R352E, or K358E substitutions displayed reductions in antiviral activities (Figure 2). rSAMD9 R304 and K358 are conserved in hSAMD9, and equivalent hSAMD9 mutations (R302E, K354E) produced similar defect in antiviral activity (Figure S2), suggesting that these residues contribute to a conserved function. However, in most mammalian SAMD9/9L proteins, the site corresponding to rSAMD9 R352 is serine (Figure 1 and S1), suggesting that rSAMD9 may rely on R352 to compensate for the missing positive charge at V354, the hSAMD9 K350-equivalent position.

### Rabbit SAMD9 targets tRNA-Phe

To further study rSAMD9 function, we established a stable HEK 293T cell line with doxycycline-inducible rSAMD9 expression. Western blotting and flow cytometry confirmed that mCherry-rSAMD9 fusion protein was expressed only upon doxycycline induction (Figure 3A and B). In the absence of rSAMD9, the cells remained permissive to vK1^-^C7^-^/GFP^+^ replication, as shown by a rising proportion of GFP-positive cells with increasing MOIs (Figure 3B&C). However, when rSAMD9 was induced, viral replication was blocked, leading to a markedly lower percentage of GFP-positive cells.

**Figure 3.**
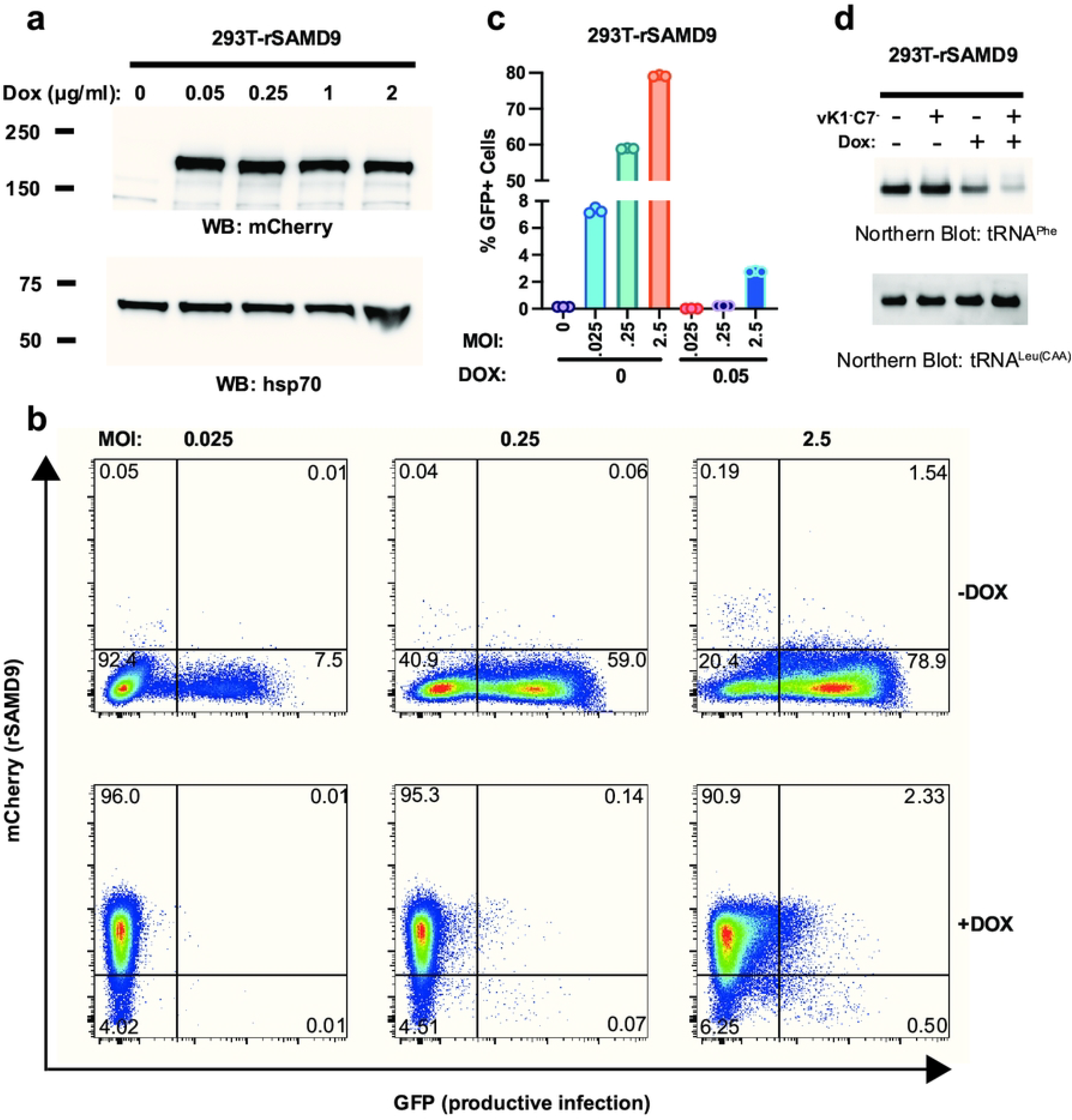
Rabbit SAMD9 specifically targets tRNA^Phe^. A stable HEK 293T cell line with doxycycline (Dox)-inducible rSAMD9 expression was established. (A). Cells were cultured with medium containing the indicated Dox concentrations. Western blotting of whole-cell lysates shows rSAMD9 (mCherry-tag) expression levels, with HSP70 as a loading control. (B). Cells were left untreated or treated with 0.05 μg/ml Dox, and then infected with vK1^-^C7^-^/GFP^+^ at the indicated multiplicities of infection (MOIs). Representative flow cytometry plots from one biological replicate are shown. (C). Quantification of the percentage of GFP+ (infected) cells from panel (B). (D). Cells were similarly untreated or treated with 0.05 μg/ml Dox and remained uninfected or were infected at an MOI of 5. RNA was harvested, and levels of tRNA^Phe^ and tRNA^Leu(CAA)^ were assessed by Northern blotting.

Since hSAMD9 restricts poxvirus replication by depleting tRNA^Phe^, tRNA^Phe^ levels were measured by Northern blot analyses following rSAMD9 induction (Figure 3D). Cells expressing rSAMD9 showed a moderate decrease in tRNA^Phe^ level, a reduction that became more pronounced after viral infection. In contrast, tRNA^Leu(CAA)^ levels were unchanged, indicating that rSAMD9, like hSAMD9, specifically targets tRNA^Phe^.

### In vitro cleavage of tRNA requires an extended tRNase module

To study the molecular basis of tRNA cleavage, we expressed and purified the recombinant rSAMD9^158-389^ protein, which corresponds to the previously characterized hSAMD9 tRNase domain. The purified rSAMD9^158-389^ protein was a monomer in solution and bound both dsDNA and tRNA^Phe^ in EMSA assays (Figure S3), consistent with earlier observations for hSAMD9 tRNase [5]. However, when incubated with either yeast tRNA^Phe^ or synthesized human tRNA^Phe^ fragment, rSAMD9^158-389^ did not produce detectable cleavage, whereas hSAMD9^134-385^ protein exhibited strong cleavage [21] (Figure 4A). Interestingly, a larger rSAMD9 fragment, rSAMD9^158-626^, which includes the SIR2 domain, was able to cleave tRNA^Phe^ in a dose dependent manner. These findings suggest that rSAMD9 requires a larger tRNase module than hSAMD9 for its nuclease activity. To determine whether the acidic residue at the site corresponding to hSAMD9 R221 was responsible for the inactivity of rSAMD9^158-389^, we also purified rSAMD9^158-389^ with an E223R substitution and tested its tRNase activity. The E223R mutant also showed negligible cleavage activities against the synthesized tRNA-Phe fragment and yeast tRNA-Phe (Figure 4B).

**Figure 4.**
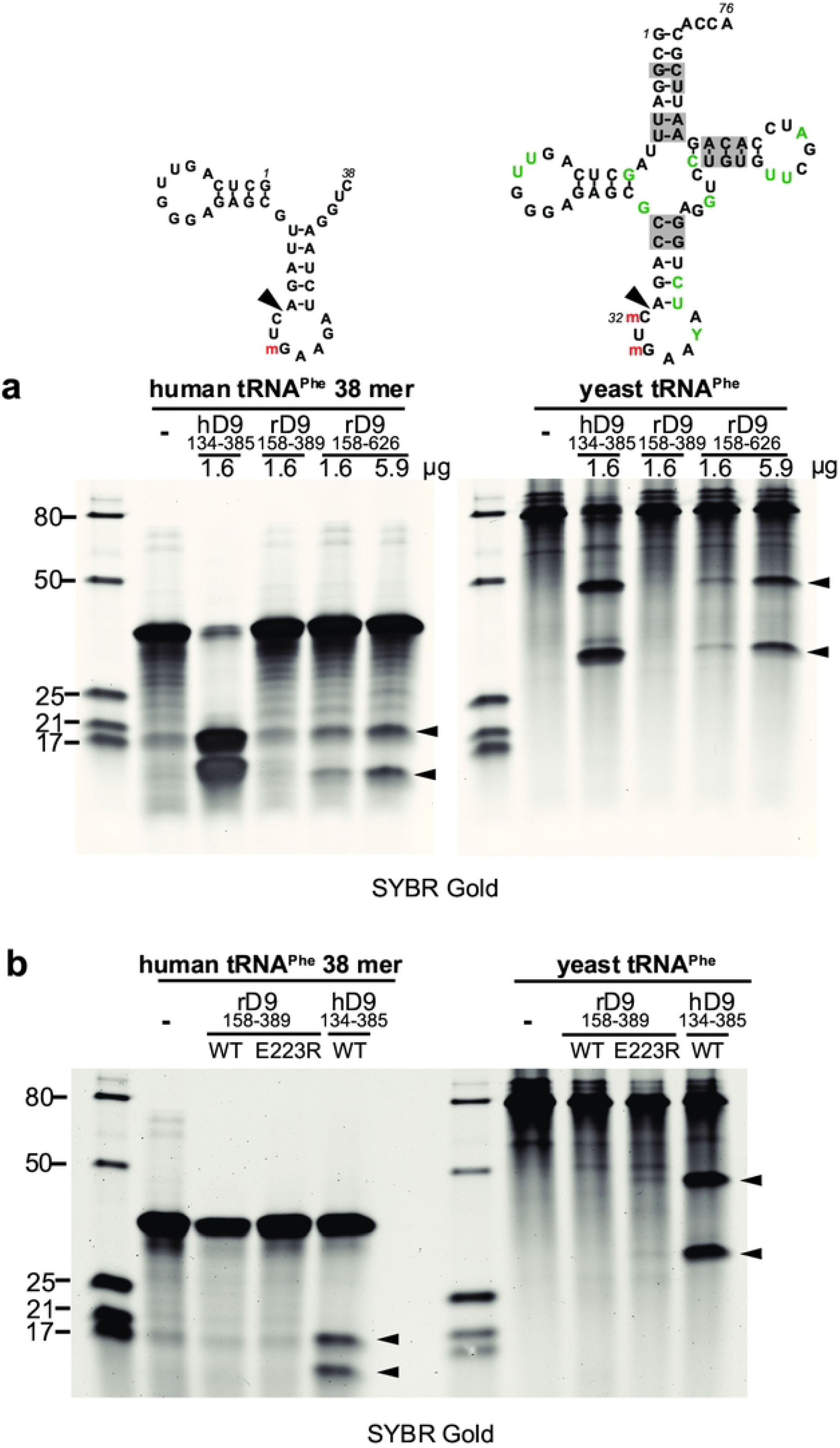
An extended rSAMD9 tRNase module is required for tRNA cleavage. A 38-nt synthetic human tRNA-Phe fragment or purified yeast tRNA was incubated for 1 hour with rSAMD9^158–389^ (WT or E22R mutant), rSAMD9^158–626^, or hSAMD9^134–385^. Cleavage products were resolved on a denaturing gel and visualized after SYBR Gold staining. The secondary structure of the RNA was depicted secondary structure was “m” indicating 2′-*O*- methylation. Sizes of the RNA ladder (in nucleotides) are indicated on the left of the gel.

### Structure analysis of rSAMD9^158-389^

Our efforts to crystalize rSAMD9^158-626^ and rSAMD9^158-389^ proteins yield suitable crystals only for the latter. The purified rSAMD9^158-389^ protein, in complex with a 26-nt dsDNA, was crystalized in space group P2_1_2_1_2, and its structure was determined to 2.4 Å resolution by molecular replacement, using hSAMD9^156-385^ structure (PDB code: 7ksp) as the search model. Four rSAMD9^158-389^ molecules occupy the asymmetric unit, forming two dimers. Overall, the rSAMD9^158-389^ structure closely resembles that of hSAMD9^156-385^, with a root-square-mean deviation (r.m.s.d.) of 0.59 Å over 178 equivalent Cα atoms (Figure 5). Nevertheless, some differences are notable. Compared with the hSAMD9 structure, the rSAMD9 structure has slightly longer β-strands at both the N- and C-termini (β1, β3, β4, β5, β6, β7, β11, and β12), as well as a short α-helix (α4) between β10 and β11. Additionally, the loop between β4 and α1 is partially disordered, missing residues 203–207, whereas the loop connecting β11 and β12 is ordered compared to the disordered loop in the hSAMD9 structure (Figure 5).

**Figure 5.**
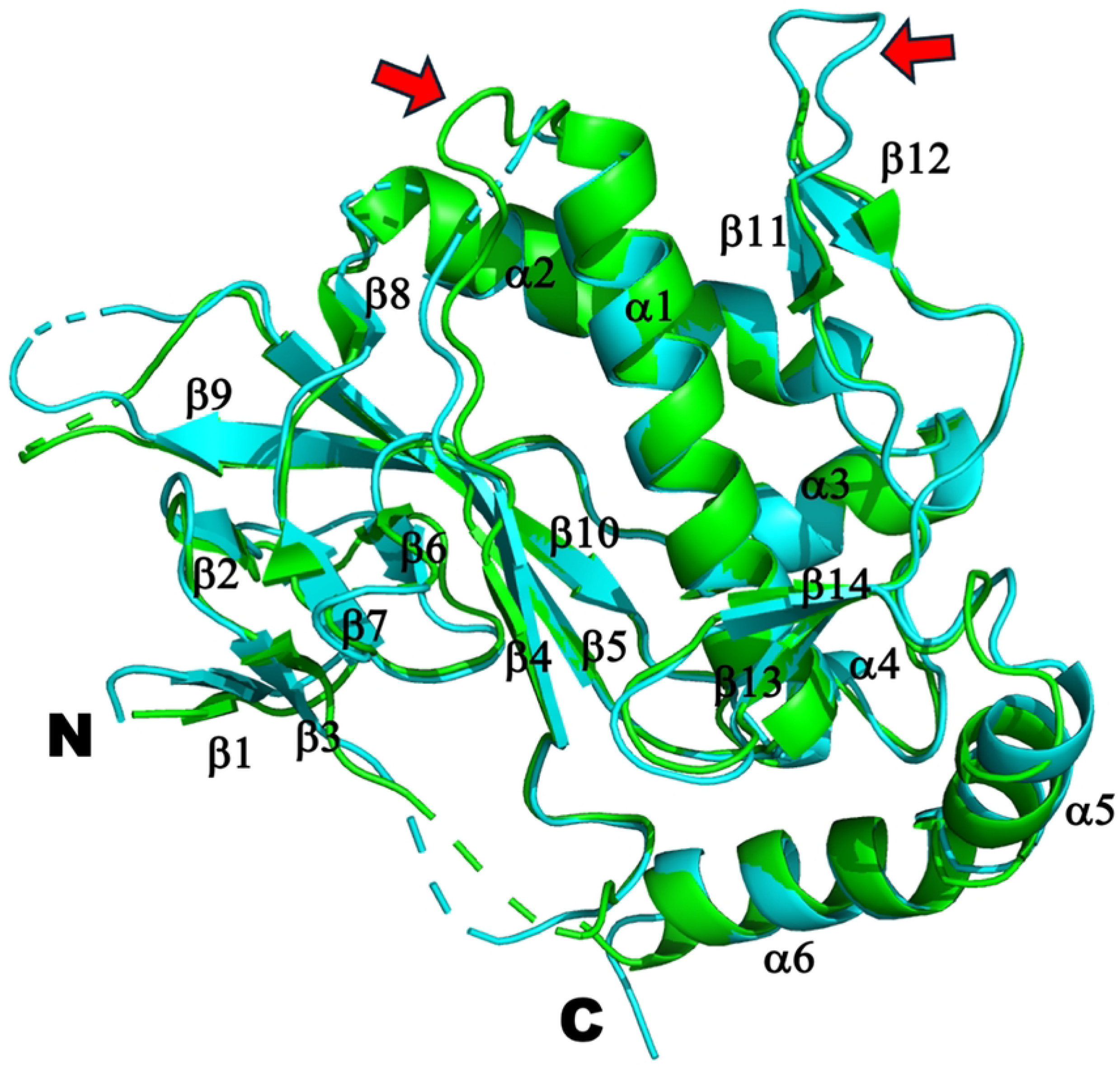
Superimposition of the structures of rSAMD9^158-389^ (cyan) and hSAMD9^156-385^ (green). The secondary structures and N- and C- termini are labeled. Red arrows indicate loops that are disordered in one of the two structures.

Surprisingly, no dsDNA densities were observed in rSAMD9^158-389^ structure, even though a stable DNA-protein complex (Fig. S3B, rSAMD9^158-389^:dsDNA at 2:1 ratio) was used for crystallization and the crystals could be strongly stained with EtBr (Figure S4). Inspection of the crystal packing revealed a space between the dimer molecules that could accommodate dsDNA in a pseudo-continuous fashion. Superimposing the rSAMD9^158-389^ dimer with the hSAMD9^156- 385^:dsDNA complex suggested that dsDNA could bind between the rSAMD9 dimers in a manner similar to hSAMD9 (Figure S5). However, the second protomer of rSAMD9^158-389^ is shifted by approximately 6 nucleotides along the dsDNA axis relative to the corresponding hSAMD9 protomer (Figure S5).

## Discussion

We recently discovered that human SAMD9 and SAMD9L function as poxvirus-activatable tRNases, regulating protein synthesis by specifically cleaving tRNA^Phe^[21]. Our structural and functional characterization of the hSAMD9 N-terminal tRNase domain revealed five positively charged residues critical for substrate binding[5,21]. Although SAMD9 and SAMD9L are paralogs that have diverged considerably, most mammalian SAMD9 and SAMD9L proteins preserve the same catalytic and tRNA-binding residues in the tRNase domain, indicating that tRNase activity is an essential, evolutionarily conserved function (Figure 1). It was therefore surprising to find that lagomorph SAMD9 orthologs harbor a negatively charged residue at one of the basic sites previously shown to be essential for hSAMD9/9L function. Through extensive structural and functional analyses of rSAMD9, we demonstrate that it retains tRNA-cleaving and antiviral properties despite notable alterations in its tRNase module, underscoring both the evolutionary conservation of tRNase activity in SAMD9/9L and the flexibility of the tRNase module within the protein family.

A key observation is that rSAMD9 remains a poxvirus-activatable tRNase (Figure 2), even though it carries an opposite charge at the hSAMD9 R221-equivalent site, a change known to inactivate hSAMD9 and hSAMD9L. We found rSAMD9 block vaccinia virus replication, whether expressed transiently or stably in cultured cells (Figures 2×3).. Furthermore, it selectively reduces tRNA^Phe^ levels in infected cells, mirroring the mechanism identified in hSAMD9 (Figure 3). Our mutagenesis studies further revealed that rSAMD9 relies on a set of key basic residues shared with hSAMD9 as well as at least one unique basic residue for full antiviral activity (Figure 2). Specifically, we confirmed the importance of two conserved basic residues (K200 and K216 in rSAMD9) that had been found in previous hSAMD9 studies, and we also identified three additional basic residues (R304, R352 and K358 in rSAMD9) in the current study. Charge-reversal mutations at rSAMD9 K200 and K216 abolished antiviral activity, while mutations of R304 and K358 caused partial defects, mirroring the effects of mutating equivalent sites in hSAMD9 (Figure S2). Moreover, our data suggest that rSAMD9 compensates for the absence of the hSAMD9 R350-equivalent residue by utilizing a species-specific basic residue (rSAMD9 R352) two amino acids upstream (Figure 1). Collectively, these findings highlight how SAMD9 can retain tRNA-binding and antiviral capacities despite deviations from certain canonical residues, emphasizing the plasticity of the tRNase binding interface.

Our crystallographic studies of rSAMD9^158–389^ supports the notion of structural conservation in the tRNase domain only with subtle variations. Comparing the hSAMD9^156–385^ and rSAMD9^158–389^ structures shows an overall similar configuration with noticeable differences in the length and flexibility of certain loops near the putative tRNA binding surface, potentially influencing function. All of the identified basic residues critical for SAMD9 function occupy the same general surface, but their relative locations in rSAMD9 and hSAMD9 structures are not perfectly superimposable (Figures 5 and 6). The absence of visible dsDNA density in rSAMD9^158–389^ crystals, despite strong ethidium bromide staining, implies that dsDNA is indeed present but may adopt multiple conformations. As previously demonstrated in hSAMD9, dsDNA binds primarily via its phosphate backbone, which may lead to flexible orientations of the nucleic acid under crystallization conditions. Thus, while rSAMD9 and hSAMD9 likely share a common substrate-binding surface, minor differences in their basic residue configuration may result in adjustments in their respective tRNase modules.

**Figure 6.**
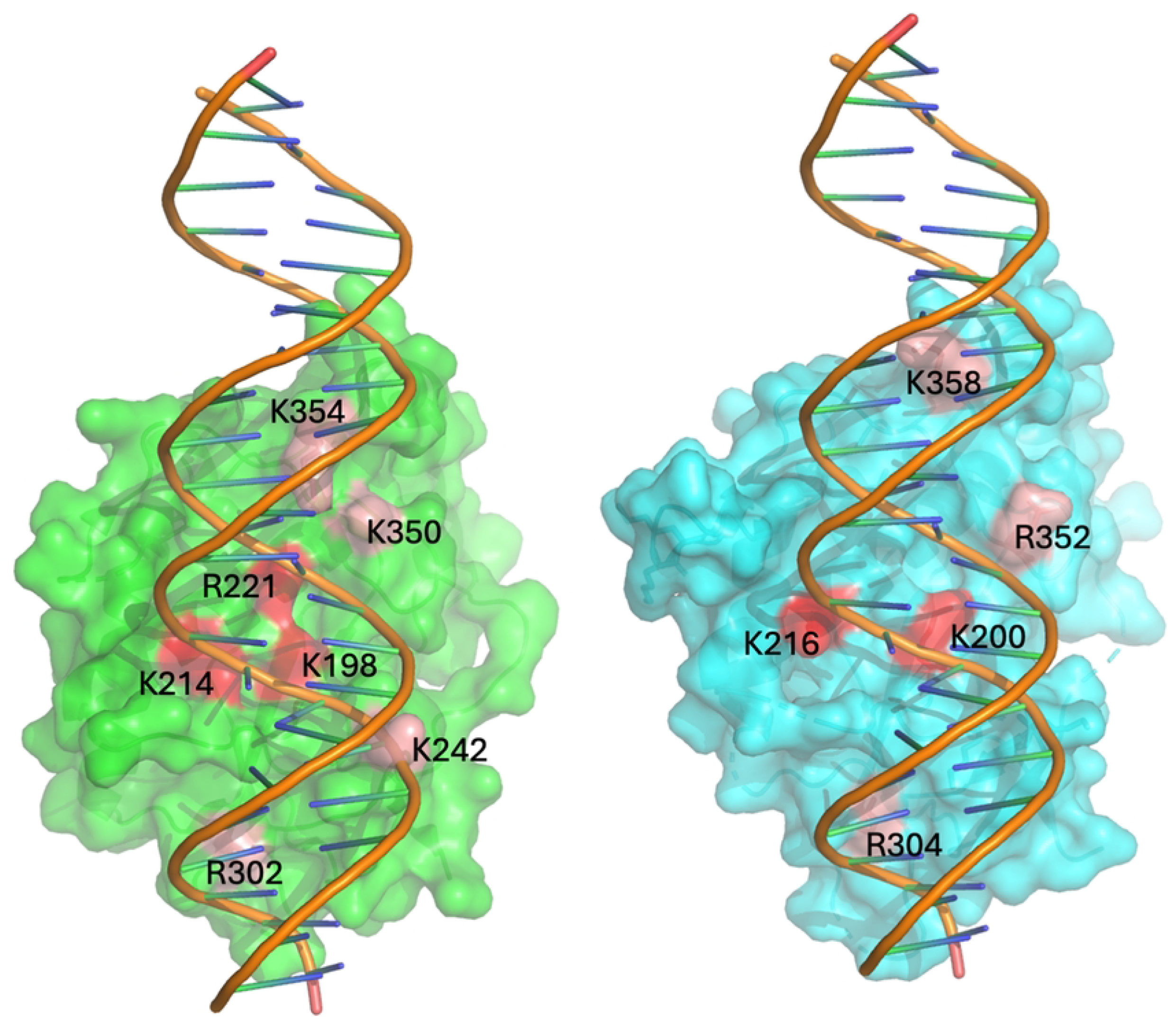
rSAMD9 and hSAMD9 share a similar substrate-binding surface. Depicted are the surface representations of the hSAMD9/dsDNA complex (left, green) and rSAMD9 with modeled dsDNA (right, cyan). The key residues for function are highlighted (red, essential; pink, important).

Another noteworthy contrast between human and rabbit SAMD9 is the size of the minimal tRNase module. Whereas hSAMD9^156-385^ alone catalyzes tRNA cleavage, the corresponding fragment in rSAMD9 (amino acids 158–389) is inactive unless extended to include the SIR2 region. The structural or functional contributions of SIR2 to tRNase activity remain unclear; it may aid substrate positioning, stabilize the active site, or influence domain dynamics. Future structural and functional studies will be needed to elucidate precisely how SIR2 regulates rSAMD9’s nuclease activity.

More broadly, our results illustrate how the SAMD9/9L family preserves a critical tRNase activity despite undergoing strong positive selection[22]. Follow-up studies can clarify whether species-specific variations affect other aspects of SAMD9 function, such as interactions with poxvirus-encoded antagonists. Finally, the spectrum of clinical disorders associated with SAMD9/9L mutations, including MIRAGE syndrome and various bone marrow failure syndromes, underscores the importance of elucidating how domain organization and specific residues modulate SAMD9/9L function. Gaining deeper insight into the antiviral, antiproliferative, and tRNA-cleaving properties of SAMD9/9L orthologs may guide novel therapeutic approaches for targeting both poxvirus infections and SAMD9/9L-related diseases.

## Materials and Methods

### Cloning rabbit SAMD9

European Rabbit SAMD9 ORF was PCR-amplified with the primer pair (5’- GGATGACGATGACAAGGCAGAACAACCGGAGCTTCCAGAAA-3’ and 5’- GGCTCCGCGGTTAAATGATCTCAATATCATAAGCAAGTGG-3’) from cDNA synthesized from RK13 cells mRNAs. A 3xFlag tag sequence was appended to the 5’ end of the ORF by PCR, and the final PCR product was cloned into pcDNA3.1 vector (Thermo Fisher Scientific). The SAMD9 ORF was completely sequenced and identical to Oryctolagus cuniculus SAMD9 reference sequence in GenBank (XP_051708952.2) except for two amino acids variations (S1008C and K1072M). All constructs were confirmed by whole plasmid sequencing (plasmidsaurus).

### Antiviral activity assay

The rSAMD9 ORF was cloned downstream of the mCherry ORF in a pcDNA6.2 vector for transient expression of mCherry and rabbit SAMD9 fusion (pcDNA6.2/mCherry-rSAMD9). Specific mutations of SAMD9 were introduced into the plasmids with Q5 site-directed mutagenesis kit (New England Biolabs, Ipswich, MA, USA). HEK 293T cells in 24-well plates were transfected with pcDNA6.2/mCherry-SAMD9 plasmids. 24 h later, the cells were infected with vK1^-^C7^-^/GFP^+^ at an MOI of 1, as previously described [5,21]. 16 h later, the cells were harvested, fixed with 4% paraformaldehyde, and analyzed with an LSR-II cell analyzer (BD Biosciences)[23]. Flow data were analyzed using FlowJo software (TreeStar).

### Inducible rSAMD9 HEK293T cell lines

A doxycycline (Dox)-inducible mCherry-rSAMD9 fusion construct was generated by subcloning into the PB-CMV-MCS-EF1-Puro vector (System Biosciences). This plasmid, along with the Super PiggyBac Transposase Expression Vector (System Bioscience), was co-transfected into HEK 293T cells. After selection, single cell clones expressing mCherry-rSAMD9 upon Dox addition were isolated following the manufacturer’s protocol.

### Northern blot analysis

Northern blots were performed as described[21]. Briefly, denaturing urea-polyacrylamide gels were used to resolve total RNA, which was then transferred onto positively charged nylon membranes. Cross-linking was carried out under short-wave ultraviolet light. DIG-labeled DNA oligonucleotide probes specific for the target tRNAs were used for hybridization, and signals were detected via chemiluminescence using CDP-Star (Roche) as the substrate.

### Protein expression and purification

European rabbit SAMD9 (residues 158-389 or 158-626) was cloned into the pXC534 vector as a 6xhis-Sumo fusion. The recombinant protein was expressed in *E. coli* BL21DE3 cells and purified in a similar way as described for human SAMD9 tRNase [5]. Briefly, the proteins were initially purified from the soluble cell lysate using a Ni-NTA affinity column. The eluted proteins then underwent Ulp1 protease cleavage, and the resultant product was collected as the flow-through of a second subtracting Ni-NTA column. Further purification to homogeneity was achieved through size exclusion chromatography. Mutants of rSAMD9^158-389^ were generated by overlapping PCR method and purified in the same way as WT. Purified proteins were concentrated to 10mg/ml, flash frozen in liquid nitrogen and store at -80°C until usage[24].

### Crystallization, structure determination and refinement

The complex of rSAMD9^158-389^ and a 26-nt dsDNA (5’ TAT TCA AAT TAA TGA TTT ATT CAA TT 3’, 5’ AT AAG TTT AAT TAC TAA ATA AGT TAA 3’) at 2:1 molar ratio was crystalized in a solution containing 0.2 M Magnesium formate dihydrate, 20% w/v Polyethylene glycol 3,350, 0.1M Hepes, pH 7.0. All data were collected at the beamline 19-ID at the Advanced Photon Source (APS), Argonne National Laboratory. The structure was solved by molecular replacement method using program Phaser[25] and the structure of hSAMD9^156-385^ (PDB code 7ksp) as the searching template. PHENIX program [26] was used for the refinement, and Coot [27] was used for the iterative manual model building. Translation, libration and screw-rotation displacement (TLS) groups used in the refinement were defined by the TLMSD server[28]. The current models are of good geometry and refinement statistics (Table S1). All molecular graphic figures were generated with PYMOL[29].

### Electrophoretic mobility shift assay

The binding of the purified rSAMD9 proteins with tRNA^Phe^ (brewer’s yeast, Sigma) was studied by electrophoretic mobility shift assay on a 0.8% native agarose gel as previously described [5].

### *In vitro* cleavage assay

Yeast tRNA^Phe^ or synthesized human tRNA^Phe^ fragment (38-mer) was incubated with 0.8 to 14 μg of recombinant rSAMD9^158–389^ protein in the cleavage buffer at 37°C for 60 min. The cleavage buffer was made of 40 mM tris-HCl (pH 8.0), 20mM KCl, 4 mM MgCl_2_ (or MnCl_2_), and 2 mM dithiothreitol(DTT). The cleavage products were mixed with gel loading dye (Invitrogen, 8546G) and resolved on 8% or 15% denaturing polyacrylamide gel containing 8 M urea. The gels were stained with SYBR Gold (Invitrogen, S11494).

## Acknowledgements

We gratefully acknowledge the staff of beam-line 19ID at the Advanced Photon Source for their support. This work was supported by NIH grant AI151638 (Y. X.). J.D. is also supported by the Oklahoma Agricultural Experiment Station at Oklahoma State University under project OKL03060 and OCAST grant HR21-071.

## Data Information

Atomic coordinates and structure factors have been deposited with the Protein Data Bank, www.rcsb.org, with accession codes 9O0D.

---

**Table 1.**
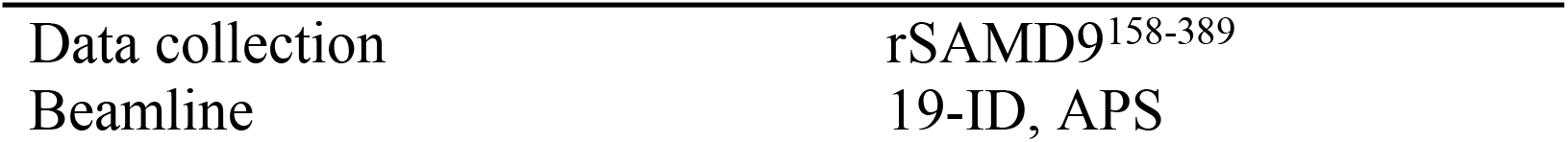
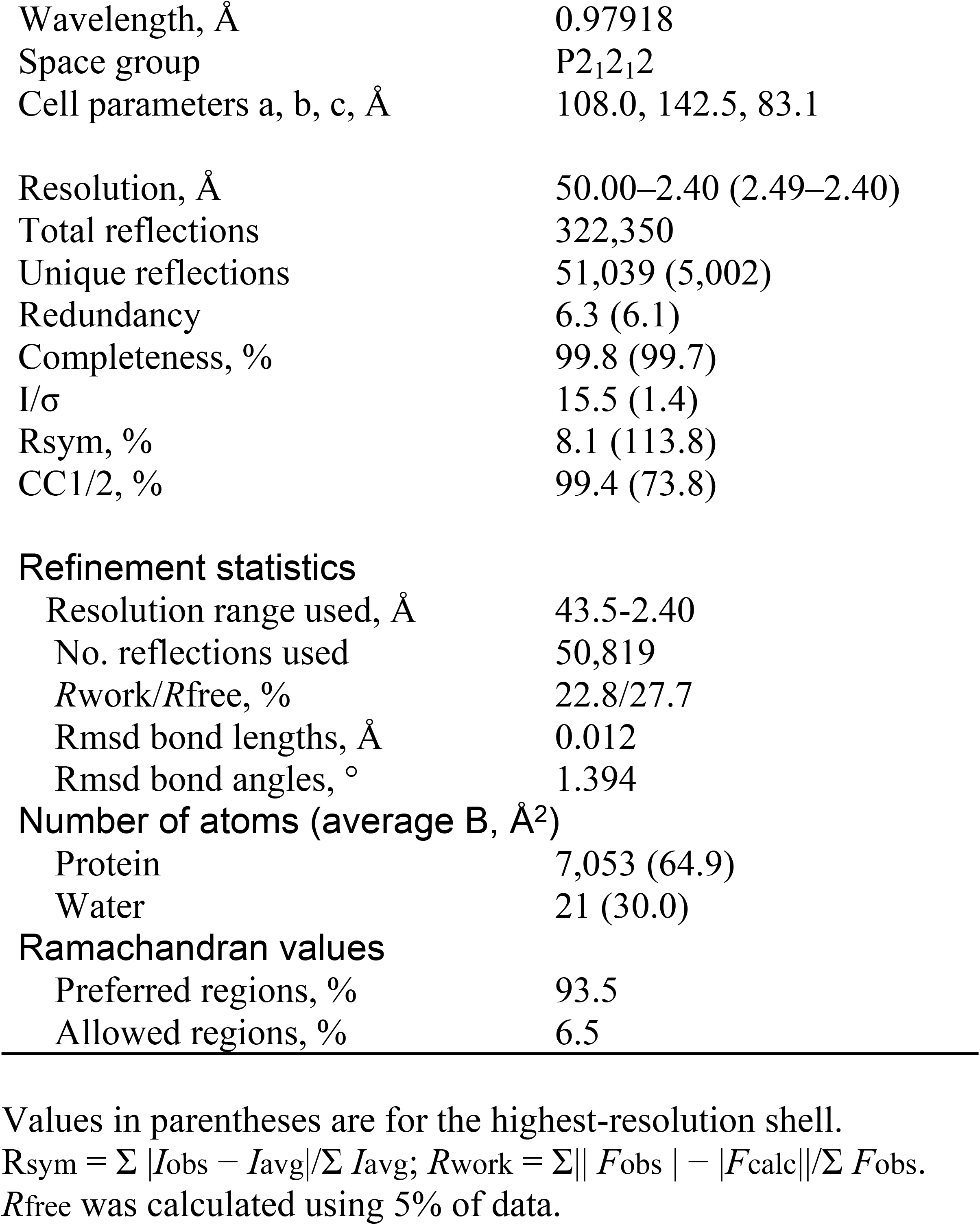
Crystallographic data and Refinement Statistics

|  |  |
| --- | --- |
| Data collection | rSAMD9<sup>158-389</sup> |
| Beamline | 19-ID, APS |
| Wavelength, Å | 0.97918 |
| Space group | P2<sub>1</sub>2<sub>1</sub>2 |
| Cell parameters a, b, c, Å | 108.0, 142.5, 83.1 |
| Resolution, Å | 50.00–2.40 (2.49–2.40) |
| Total reflections | 322,350 |
| Unique reflections | 51,039 (5,002) |
| Redundancy | 6.3 (6.1) |
| Completeness, % | 99.8 (99.7) |
| I/σ | 15.5 (1.4) |
| Rsym, % | 8.1 (113.8) |
| CC1/2, % | 99.4 (73.8) |
| Refinement statistics |  |
| Resolution range used, Å | 43.5–2.40 |
| No. reflections used | 50,819 |
| *R*work/*R*free, % | 22.8/27.7 |
| Rmsd bond lengths, Å | 0.012 |
| Rmsd bond angles, ° | 1.394 |
| Number of atoms (average B, Å<sup>2</sup>) |  |
| Protein | 7,053 (64.9) |
| Water | 21 (30.0) |
| Ramachandran values |  |
| Preferred regions, % | 93.5 |
| Allowed regions, % | 6.5 |

Values in parentheses are for the highest-resolution shell.
$R_{\text{sym}} = \Sigma |I_{\text{obs}} - I_{\text{avg}}| / \Sigma I_{\text{avg}}$ ; $R_{\text{work}} = \Sigma ||F_{\text{obs}}| - |F_{\text{calc}}|| / \Sigma |F_{\text{obs}}|$ .
*R*free was calculated using 5% of data.

